# Genome-wide analysis of the association of transposable elements with gene regulation suggests that *Alu* elements have the largest overall regulatory impact

**DOI:** 10.1101/142018

**Authors:** Lu Zeng, Stephen M. Pederson, Danfeng Cao, Zhipeng Qu, Zhiqiang Hu, David L. Adelson, Chaochun Wei

**Affiliations:** School of Life Sciences and Biotechnology Shanghai Jiao Tong University Shanghai P. R. China; School of Biological Sciences The University of Adelaide Adelaide, SA Australia

**Keywords:** Transposable elements, Epigenetic, Gene expression, Gene ontology, Regulatory elements

## Abstract

Nearly half of the human genome is made up of transposable elements (TEs) and there is evidence that TEs are involved in gene regulation. Here, we have integrated publicly available genomic, epigenetic and transcriptomic data to investigate this in a genome-wide manner. A bootstrapping statistical method was applied to minimize the confounder effects from different repeat types. Our results show that although most TE classes are primarily associated with reduced gene expression, *Alu* elements are associated with up regulated gene expression. Furthermore, *Alu* elements had the highest probability of any TE class of contributing to regulatory regions of any type defined by chromatin state. This suggests a general model where clade specific SINEs may contribute more to gene regulation than ancient/ancestral TEs. Finally, non-coding regions were found to have a high probability of TE content within regulatory sequences, most notably in repressors. Our exhaustive analysis has extended and updated our understanding of TEs in terms of their global impact on gene regulation, and suggests that the most recently derived types of TEs, *i.e.* clade or species specific SINES, have the greatest overall impact on gene regulation.

## INTRODUCTION

Repetitive elements are similar or identical DNA sequences present in multiple copies throughout the genome. The majority of the repetitive sequences in the human genome are derived from transposable elements (TEs) (Lander, et al. 2001; Smit 1999) that can move within the genome, potentially giving rise to mutations or altering genome size and structure. Typical eukaryotic genomes contain millions of copies of transposable elements (TEs) and other repetitive sequences. TEs fall into two major classes: those moving/replicating via a copy and paste mechanism and an RNA intermediate (retrotransposons) and those moving via direct cut and paste of their DNA sequences (DNA transposons). Retrotransposons can be subdivided into two groups: Those with long terminal repeats (LTRs), and those without LTRs (non-LTRs). Human LTR elements are related to endogenous retroviruses (HERVs), which along with similar elements account for nearly 8% of the human genome (Cordaux and Batzer 2009). Non-LTR retrotransposons include two sub-types: autonomous long interspersed elements (LINEs) and non-autonomous short interspersed elements (SINEs), which are dependent on autonomous elements for their replication; both LINEs and SINEs are widespread in eukaryotic genomes. LINE-1 (long interspersed element 1) and *Alu* elements are two TEs that belong to non-LTR retrotransposons, which account for approximately one-quarter of the human genome (Lander, et al. 2001).

A number of existing studies have shown that TEs can influence host genes by providing novel promoters, splice sites or post-transcriptional modification to re-wire different developmental regulatory and transcriptional networks (Cowley and Oakey 2013; Kunarso, et al. 2010; Lynch, et al. 2011). TEs tend to regulate gene expression through several mechanisms (Britten 1996; Cowley and Oakey 2013; Pereira, et al. 2009; van de Lagemaat, et al. 2003). For example, in rodents, the expression levels of protein coding genes containing repetitive elements are significantly associated with the number of repetitive elements in those genes (Pereira, et al. 2009). L1 elements show a stronger negative correlation with expression levels than the gene length (Jjingo, et al. 2011), and the presence of L1 sequences within genes can lower transcriptional activity (Han, et al. 2004). Moreover, TEs have been shown to influence gene expression through non-coding RNAs, resulting in the reduction or silencing of gene expression (Rebollo, et al. 2012). For example, the expression of long intergenic non-coding RNAs (lincRNAs) was strongly correlated with HERVH transcriptional regulatory signals (Kelley and Rinn 2012). Past studies have found that TEs have contributed to nearly half of the active regulatory elements of the human genome (Jacques, et al. 2013), by altering gene promoters and creating alternative promoters and enhancers to regulate gene activity (Conley, et al. 2008; Franchini, et al. 2011; Medstrand, et al. 2001). According to previous research, 60% of TEs in both human and mouse were located in intronic regions and all TE families in human and mouse can exonize, supporting the view that TEs may create new genes and exons by promoting the formation of novel or alternative transcripts (Piriyapongsa, et al. 2007; Sela, et al. 2007). The association between repetitive elements and RNAs has also been investigated. For example, *Alu* elements in lncRNAs can lead to *STAU1* mediated mRNA decay by duplexing with complementary *Alu* elements in the 3’UTRs of mRNAs (Hadjiargyrou and Delihas 2013), and the insertion of TEs may also drive the evolution of lincRNAs and alter their biological functions (Kelley and Rinn 2012). In this study, TEs in the human genome were analyzed using genome-wide datasets associated with gene regulation. These datasets enabled an assessment of the association of TEs with Regulatory Elements (RE) based on chromatin states, as marked by histone modification within six human cell lines, lincRNA, Gene Ontology (GO) enrichment, as well as overall transcriptome profiles.

Although, a number of studies have analyzed the association between TEs and gene regulation in the human genome, they did not consider whether gene intervals contained multiple repeat types or not. This can cause a confounding factor when attempting to determine the regulatory impact of a specific repeat type. In our study, we eliminated confounding effects by focusing on gene intervals that only contained one type of repeat and used a weighted bootstrap approach to minimize any influence of co-occurring elements. Our method presents a novel way to investigate the association of specific TE type with gene expression.

## RESULTS

### The Distribution Of Repetitive Elements In The Human Genome

Initially, we compared the distribution of repetitive elements in the human genome. We found many repetitive elements overlapped with gene models from the human RefSeq gene datasets, and their distributions with respect to components of gene models are shown in Figure 1. Fewer repetitive elements were found in coding exons and non-coding exons such as the 5’ and 3’ UTRs. This indicates that repeat accumulation in these regions is selected against in order to preserve regulatory and coding functions (Figure 1).

**Fig. 1.**
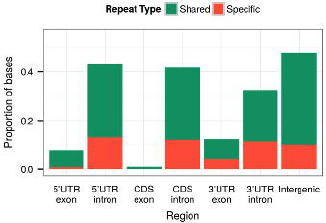
Distribution of repetitive elements overlapping with different human gene regions. Gene regions are shown on the x-axis and the y-axis shows the percentages of the genomic regions containing repetitive elements. Human-specific repeats were those annotated with “Homo sapiens” or “primates” as their origin (see Table S8 for the list of human-specific repeat classes). The remaining repetitive regions were categorized as shared repeats.

### Transposable Elements And Gene Regulation By Chromatin States

It is known that TE-derived sequences can contribute to transcription factor binding sites, promoters and enhancers, and insulators/silencers (de Souza, et al. 2013; Lynch, et al. 2011). To look for enrichment of TEs within regulatory elements, we looked at the proportions of nucleotides with a TE in six defined regulatory elements (Ernst, et al. 2011) as they appear in different components of the gene model. This represents the probability of a given nucleotide within a Regulatory Element (RE) being from a transposable element (TE), i.e. p(TE|RE) (see Methods for details) across the set of genic regions (Figure 2A). One major observation from Figure 2A is that TE have higher p(TE|RE) in insulators indicating that they are present at higher levels in insulators relative to most other regulatory regions. While Active Promoters have the lowest repeat level in regulatory regions. The proportion of TEs in RE is lower than the genome wide average based on the p(TE|RE) values with exceptions being most Weak Enhancers in non-coding regions. Unsurprisingly, CDS exons have the lowest repeat content. The overall under-representation of TE in RE is likely indicative of negative selection for TE in regulatory regions.

**Fig.2.**
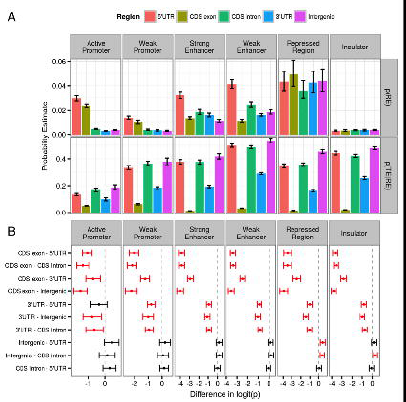
Analysis of the co-occurrence of Transposable Elements and Regulatory Elements across multiple genomic regions. A) Probability estimates are those of an individual base within each region being part of a Regulatory Element, i.e. p(RE), or a Transposable Element within a Regulatory Element, i.e. p(TE|RE). Error bars indicate ±1 Std. Error as calculated on the logit-transformed values. B) Pairwise comparisons of TE content in RE based on 1 – *α* Confidence Intervals for the difference between logit-transformed probabilities p(TE|RE), adjusted for multiple comparisons at the level *α* = 0.05/6 within each RE (60). Intervals highlighted in red show significant pairwise differences (confidence intervals do not cross the 0 difference value).

Pairwise comparisons of p(TE|RE) between different gene model compartments are shown in Figure 2B in order to identify significant differences (see methods for description of statistical methods used) in repeat content between regulatory regions/compartments. These reveal that for all regulatory elements, TEs are more sparsely distributed across regulatory elements within CDS exons than regulatory elements in other genic regions. Likewise, regulatory elements in the 3’UTR were more sparsely populated with TEs in comparison to those in other regions, with the sole exception being Active Promoters in the 5’UTR. In contrast, Polycomb Repressed Regions and Insulators in intergenic regions were enriched for TEs in comparison to these elements in other gene model compartments.

### Different Classes Of Transposable Elements And Their Associations With Chromatin State

In order to systematically characterize the potential role of different repeat classes within the regulatory elements mentioned above, the distribution of regulatory elements within specific TEs classes were investigated as below, using the probability of a nucleotide also belonging to a TE element p(TE|RE) (See Methods for details). Six TE classes (*Alu*, L1, L2, LTR, MIR and DNA transposons) were examined (Figure 3A). The most significant observation from Figure 3A was that *Alu* had the highest p(TE|RE) across all regulatory regions, especially in Weak Promoter, Strong Enhancer and Weak Enhancer, while recent inserted L1 elements were least likely to be within regulatory elements compared to the other five repeat classes. Although L1 elements are the most abundant mobile elements in the human genome, accounting for 17% of the human genome, they had lower p (TE|RE) compared to *Alu* elements, which may indicate a negative selection for L1 element insertions in gene regulatory regions. Interestingly, MIR and L2, ancient repeats that have been inactive since prior to the mammalian radiation, are present at very low levels in the human genome and they have a relatively high p(TE|RE) in Strong Enhancer regions.

**Fig.3.**
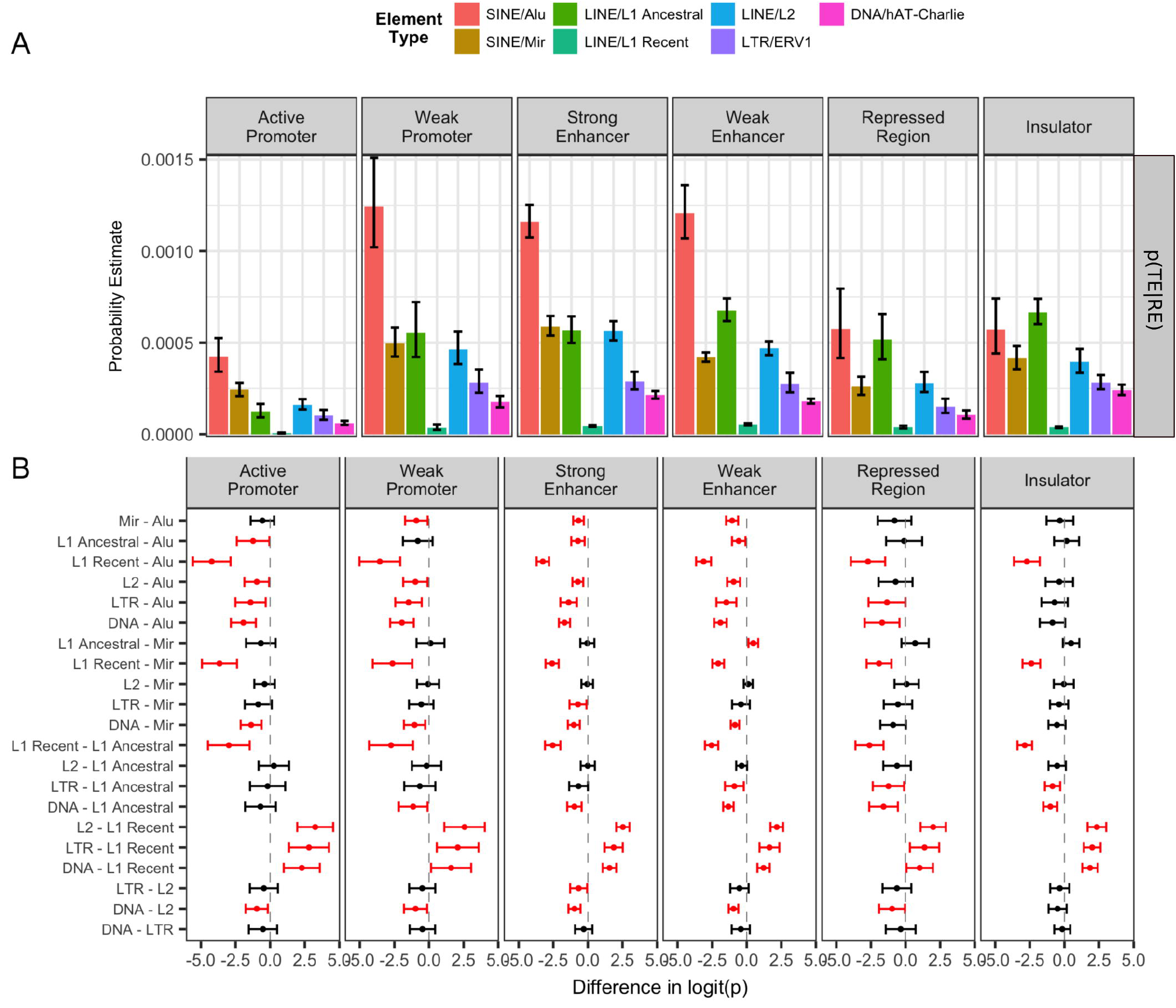
Analysis of the occurrence of Regulatory Elements within specific classes of Transposable Elements. A) Probability estimates are for an individual base within each type of element belonging to each of the regulatory elements, i.e. p(RE|TE). Error bars indicate ±1 Std. Error as calculated on the logit-transformed values. B) Pairwise comparisons of TE content in RE based on 1*–α* Confidence Intervals for the difference between logit-transformed probabilities p(RE|TE), adjusted for multiple comparisons at the level *α* = 0.05/6 within each RE (60). Intervals highlighted in red show significant pairwise differences (confidence intervals do not cross the 0 difference value).

In order to identify significant differences within specific repeat types between regulatory regions, pair-wise comparisons of p(TE|RE) were implemented and shown in Figure 3B (see methods for description of statistical methods used). Among six defined regulatory regions, *Alu* elements contributed more to Active Promoter, Weak Promoter, Strong Enhancer and Weak Enhancer regions compared to the other five repeat types. Ancestral L2 and MIR repeats were found to be retained more than recent L1 across all RE. Another indication of potential exaptation of ancestral repeats was the observation that recent inserted L1 had the lowest probability of being in a RE compared to other repeat types (Figure 3B). However, LTR elements and DNA transposons were also more likely to be retained than recent L1s, suggesting some degree of exaptation for those repeats as well.

### TEs Are Abundantly Present In Long Intergenic Non-Coding RNAs In Regulatory Elements

Because ncRNAs are known to regulate gene expression, we decided to examine the relationships between lincRNAs and TEs in REs (Figure 4). The observation that >30% of nucleotides from many of the regulatory elements were derived from TEs was quite striking. Specifically, lincRNA exonic regions contained the highest RE density for Polycomb Repressed Regions (Figure 4A), with nearly a third of these nucleotides being derived from TEs, suggesting that the presence of transposable elements in lincRNA exons may be associated with regulation of lincRNA expression and subsequently linked to gene regulation. Unsurprisingly, we found that TEs made up a lower proportion of nucleotides in CDS-exons across all regulatory elements, when compared to CDS-introns, lincRNA exons and lincRNA introns (Figure 4B).

**Fig.4.**
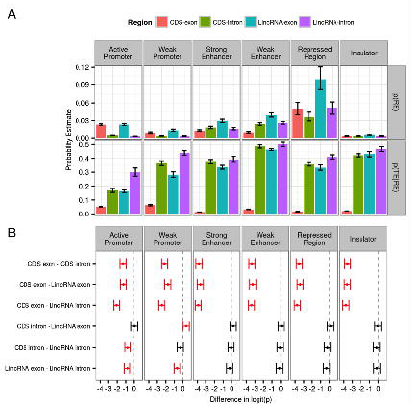
Analysis of the co-occurrence of Transposable Elements and Regulatory Elements across non-coding regions. A) Probability estimates for an individual base within each type of non-coding region being part of a Regulatory Element p(RE) or a Transposable Element within each Regulatory Element p(TE|RE). Error bars indicate ±1 Std. Error as calculated on the logit-transformed values. B) Pairwise comparisons of TE content in RE based on 1*–α* Confidence Intervals for the difference between logit-transformed probabilities p(TE|RE), adjusted for multiple comparisons at the level *α =*0.05/6 within each RE (60). Intervals highlighted in red show significant pairwise differences (confidence intervals do not cross the 0 difference value).

### Identification of biases due to multiple TE types and transcript length on gene expression

We first summarized the overall distribution of transposable elements within various compartments of gene models by finding genes containing TEs in either single or multiple compartments (Figure S1, Table 1), and genes containing one or more type of TE (Figure S2, Table 2). Because gene/transcript length can also cause bias in the estimation of gene expression we examined the relationship between gene length and which compartments of a gene contained a TE (Figure S3), as well as the relationship between gene length and the presence of a specific type of TE (Figure S4), using a Wilcoxon Test (Tables S2 & S3). We found that only genes with TEs in the 3’UTR or in multiple genic regions showed a bias towards longer length, whilst for TEs exclusively within the proximal promoter or 5’UTR there was a bias towards shorter genes (Figure S3; Table S2). When assessing the relationship between gene length and the presence of a specific TE class, the length of genes with *Alu*, L2 or MIR elements alone were very similar to genes with no TE, whilst L1 and LTR elements showed a bias towards shorter genes, and the presence of multiple elements biased towards longer genes (Figure S4; Table S3).

### Effects On The Probability Of A Gene Being Detected As Expressed Due To The Presence Of A TE Across The Different Gene Model Compartments

As chromatin states are not always indicative of changes in transcriptional activity, we further investigated any effects on human gene expression due to the presence of specific TE classes within each of the four regulatory regions, i.e., Proximal Promoter, 5’UTR, CDS and 3’UTR. However, as TEs are far less frequent in CDS regulatory regions (Figure S1), the subsequent analysis focused on the other three regions. Six human tissue transcriptome datasets (adipose, brain, kidney, liver, skeletal muscle and testes tissue) were selected from the Illumina BodyMap2 dataset for this analysis, and global patterns of gene expression were investigated based on the presence or absence of each TE within these three genic regions respectively.

In order to exclude confounder effects caused by multiple repeat types on the expression of single genes, the weighted bootstrap method was applied to both the probability of a gene being detected as expressed (Figure 5A), and to the overall expression levels for those genes detected as expressed (Figure 5B). This revealed that *Alu* elements are commonly associated with a higher probability of expression when located in either the 5’UTR or the 3’UTR across the majority of tissues. In contrast to the presence of an *Alu*, the presence of L1 elements in the Proximal Promoter showed a negative impact on the probability of a gene being detected as expressed in 3 out of the 6 tissues, with the remaining tissues being directionally consistent and quite likely to be Type II errors (Supplementary Figure S6A).

**Fig.5.**
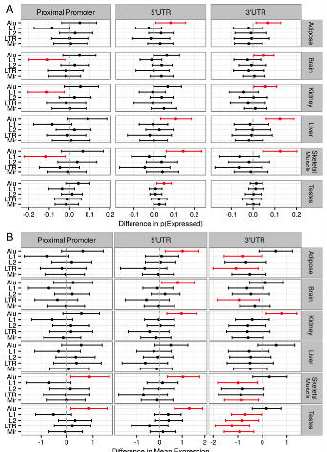
Effects on gene expression of the presence of specific TEs in each gene compartment. A) Confidence Intervals for the difference in the probability of a gene being detected as expressed due to the presence of each TE in each genic region. B) Confidence Intervals for the difference in mean log2(TPM) counts. For both A) and B), Confidence Intervals were obtained using the weighted bootstrap and are **1–*α/m*** intervals, where ***α*=0.05** and ***m*=90** as the total number of intervals presented. Dots represent the median value from the bootstrap procedure, whilst the vertical line indicates zero. Intervals which do not cross zero are coloured red, and indicate a rejection of the null hypothesis, **H_0_:*Δθ* = 0**, where *θ* represents the parameter of interest.

### Effects On The Levels Of Gene Expression Due To The Presence Of A TE In Each Gene Model Compartment

Again using the weighted bootstrap approach to minimize any confounding effects of co-occurring elements, the presence of an *Alu* in the 5’UTR was found to be associated with increased expression levels in five of the six tissues investigated (Figure 5B). Similarly, *Alu* elements in the Proximal Promoter were associated with increased expression in two of the tissues (skeletal muscle and testis). *Alu* elements in 3’UTR were associated with elevated expression levels only in the Kidney sample. The presence of L1, L2 and MIR elements showed varying degrees of reduced gene expression across the tissues when located in the 3’UTR only. It was also noted that whilst strongly controlling the family-wise Type-I error rate (FWER), the adjusted confidence intervals would result in an increase in the Type-II error rate where true differences cannot be detected. As such, the point at which the confidence intervals would include zero was found and taken as a proxy for the p-value. Confidence intervals based on these p values to an FDR of 0.05 are shown in Supplementary Figure S6 with the p values given in Supplementary Table S5. It is clear from this additional approach that the role of TEs such as L1 elements in Proximal Promoters and 3’UTRs, LTR elements in 5’UTRs and many of the elements in the 3’UTR may have been considerably understated as a result of our more conservative approach.

### Analysis Of Genes With Exapted Or Exonized TEs

In order to determine if exapted or exonized TEs might influence gene function, genic regions (proximal promoter, 5’UTR and 3’UTR) that overlapped with TEs were analyzed to assess the association of TEs with different gene functions (annotated using Gene Ontology).

The three fundamental GO categories are: cellular component, molecular function and biological process. Enrichment information for each GO category is listed in Supplementary Table S6. First, we looked at the association of gene function and repeats in the biological process category (Figure 6, Table S6a). Figure 6 represents that 5’UTR regions containing repeats showed a strong enrichment in genes with regulatory processes, mainly involving metabolic processes, while promoter regions containing repeats were rarely associated with biological process. A similar pattern was found with respect to cellular component (Figure S7, Table S6b) and molecular function (Figure S8, Table S6b), repeats in 5’UTR were predominantly enriched in intracellular/cytoplasmic structures and associated with binding activities respectively.

**Fig.6.**
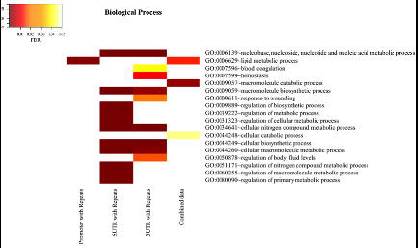
Enrichment of GO terms of genes containing TEs in promoter, 5’UTR and 3’UTR regions. Enrichment of GO terms of genes containing TEs in “Biological Process”. Genes containing different types of repetitive elements in the proximal promoter regions are labeled as "Promoter with Repeats", and Genes containing repetitive elements in UTR regions are labeled as "5/3UTR with Repeats". Genes named “Combined Repeats” are the combined data from 3 regions we mentioned above. The darker the color, the greater the GO term enrichment as determined by FDR.

The same method was applied to investigate the association of repeats with function in protein coding regions (CDS). We found that genes with protein coding exons containing Alus were enriched for the GO term “intracellular non-membrane-bounded organelle”. Interestingly, these exonization/exaptation events were found associated with splice variants incorporating *Alu* sequences (Table S7 & Figure S9, S10).

Moreover, according to our analysis of TEs and alternative splicing data, we found that 2.98% of alternatively spliced transcripts contained TEs within protein coding exons (Table 3). *Alu* and MIR were the two most common repeats to be involved in alternative splicing/exonization, but LINEs and LTR also contributed to a significant proportion of exonization. This suggests a role for exonization of TEs in the evolution of increasing coding and regulatory versatility of the transcriptome.

## DISCUSSION

In this work, we analyzed the distribution of various classes of transposable elements and their association with regulatory elements (active chromatin) for the first time. We have also analyzed the association of specific repeat types with gene expression in the human genome using a new method that minimizes confounders. Based on the analysis of the TE distributions in genic regions and corresponding gene expression patterns, the presence of specific TEs was found to be associated with changes in gene expression. Further, gene function as defined by GO term analysis differed depending on the TE insertion site within the gene. Finally, we looked at TEs present in ncRNAs, specifically lincRNAs, and found that repetitive elements were present at higher levels in lincRNAs than in coding exons.

### TEs And Gene Regulation

It is known that epigenetic modification of TEs can regulate transcription by altering active chromatin, and this chromatin modification has evolved from epigenetic ‘defense’ mechanisms to silence TE activity (Slotkin and Martienssen 2007). Here we present a comprehensive analysis of TE contribution to each gene compartment associated with different types of active chromatin. A role for TEs in silencing is consistent with our finding that they are enriched in Poly-comb repressed regions and Insulators (Figure 2). However, the enrichment of TEs in Weak Enhancer and Strong Enhancer regions (Figure 2) is suggestive of exaptation of TEs as active regulators of gene transcription. We also found that regulatory regions overlapping with genes had a greater proportion of sequence originating from TEs in the 5’ and 3’UTRs compared to coding exons (Fig 2). This is not surprising considering the potential adverse effect of TE insertion in a protein coding sequence (Belancio, et al. 2006; Belancio, et al. 2008).

TEs have been implicated in altering gene expression through a number of mechanisms, for example, TE can be exonized through alternative splicing and they can alter transcription factor binding because they contain transcription factor binding sites (Cordaux and Batzer 2009; Polak and Domany 2006). However, it is difficult to analyze the impact of specific TEs on gene expression, due to the co-occurrence of other TE types in or near individual genes. To overcome this problem, we used a bootstrapping approach for gene expression analysis, effectively allowing us to focus on genes containing single TE types. Figures 5B and S6B summarize our findings with regard to gene expression, and indicate that *Alu* elements in proximal promoter regions, 5’UTR and 3’UTR are commonly associated with increased gene expression. Moreover, *Alu* elements were found to have a high probability of contributing to regulatory elements, especially with respect to Weak Promoter, Strong Enhancer and Weak Promoter regions (Figure 3). These results are consistent with previous reports that motifs in TEs such as Alus can be exapted as transcription factor binding sites and are involved in gene regulation (Kim, et al. 2004; Levanon, et al. 2004; Lin, et al. 2009). However, in contrast to our finding that *Alu* elements were associated with increased gene expression, the exonization of *Alu* sequences from alternative splicing seems to cause only decreased expression of the alternatively spliced transcript (Lin, et al. 2009). Furthermore, the persistence of *Alu* insertions would most likely be selected against, which would suggest that an *Alu* element can rarely take on a regulatory role when it is inserted into a gene (Deininger 2011).

Although L1 elements are the most abundant repeats in the human genome, they were associated with decreased gene expression (Fig 5), and were less prevalent in regulatory elements or active chromatin, when compared to other repeat classes (Fig 3). Considering the size of L1 elements (~6kb), it is easy to understand that the insertion of L1 elements, even if truncated, are likely to cause disruption of host gene function (Kazazian, et al. 1988; Meischl, et al. 2000; Perepelitsa-Belancio and Deininger 2003). Therefore, newly inserted L1 elements are probably under negative selection in regulatory regions and are less likely be exapted as a RE.

L2 and MIR are two ancient TE families conserved in mammals, and are inactive or fossil TE elements (Deininger and Batzer 2002). According to previous studies MIR elements have been exapted as functional enhancers in the human genome, and are enriched in transcription factor binding sites (Jjingo, et al. 2014). However, our results showed that MIR and L2 elements were associated with reduced gene expression when located in 3’UTRs (Figure 5), but not with increased gene expression. This indicates that their exaptation as enhancers has the opposite effect of their exaptation within gene models. Figure 5 also showed that LTRs were associated with repression of gene expression, which is in contrast to previous work that implicated LTRs as alternative promoters (Cohen, et al. 2009). ERVL LTRs have been reported to provide molecular mechanisms for stochastic rewiring of gene expression and evolution in the mammalian germline (Franke, et al. 2017). Our analysis suggests that this rewiring does not affect the majority of expressed genes except for LTR inserted into 3’UTR, which are associated with decreased gene expression.

### The Association Between TEs And LincRNA

LincRNAs are known to regulate gene expression through epigenetic mechanisms (Engreitz, et al. 2013) and can be important regulators of the human innate immune response (NE, et al. 2014). One possible factor driving lincRNA evolution and biological function is the insertion of TE elements, TE insertions into lincRNA were confirmed significantly reduced expression in many tissues and cell lines than lincRNAs that are devoid with TEs (Kelley and Rinn 2012). This transcriptional repression can be supported with our results, as we found lincRNA containing TEs were clearly enriched in Insulators and Polycomb repressed regions (Figure4). However, we also found that lincRNA containing TEs were highly enriched in Weak Enhancer regions (Figure 4), suggesting that TEs present in lincRNA may have an ‘active’ role in gene expression.

### The Exaptation of TEs

In addition to potentially altering gene expression by insertion into regulatory elements, TEs may also be associated with specific functional characteristics of expressed protein coding genes. When we examined the functional annotation of repeat containing genes, we found that some functions were over-represented (Table S5). Perhaps the most interesting of these associations was that genes with *Alu* insertions were found to contribute to coding exons through alternative splice variants. One explanation of this observation is that *Alu*-induced alternative transcripts may result in nonsense-mediated decay of alternative transcripts (Lewis, et al. 2003). Two examples of alternatively spliced genes of this type with implications for human disease are *DISC1* and *NOS3* (Table S7 and Figure S9 & S10). *DISC1* alternative transcripts are known to contribute to increased risk of schizophrenia (Callicott, et al. 2005; Rapoport, et al. 2005) and *NOS3* transcript variants are associated with cardiovascular disease phenotypes (Hingorani, et al. 1999; Pacanowski, et al. 2009). Based on previous research, nearly 4% of protein-coding sequences include transposable elements, and one-third of them are *Alu* insertions (Nekrutenko and Li 2001). Therefore, *Alu* exonization in protein-coding genes may play an important role in modifying gene expression.

### Implications For Our Perception Of Genome Evolution

While there are many publications implicating TEs in the regulation of individual genes, our work clarifies some previous uncertainties and resolves some contradictions, confirming that this role of TEs is significant across the genome. In general, most TEs would appear to be strongly associated with repression of gene expression, either through the 5’UTR or perhaps as components of lincRNA exons. However, the presence of Alus in 3’UTR and proximal promoter regions may act to increase gene expression. These results are supported by previous studies (Jjingo, et al. 2011) and provide a new understanding of how repeats are associated with epigenetic regulation of gene expression. Finally, while exapted TEs may contribute to the generation of transcripts that undergo nonsense mediated decay as part of gene regulation, we speculate that they may also provide an opportunity for alternative splicing and novel exaptation. TEs therefore are important agents of change with respect to the evolution of gene expression networks.

## MATERIAL AND METHODS

### Theoretical Framework And Methods

We constructed pipelines to analyze the distribution of repetitive elements in different parts of the human genome. Repetitive elements overlapping with protein coding regions, non-coding regions and regulatory elements were identified. GO term over-representation and expression analyses were carried out for repetitive elements overlapping with protein-coding regions. The pipelines and related materials are described below.

### Tools Used To Develop Pipelines For Repetitive Element Analysis

The identification and classification of TEs from the human genome was conducted by developing a pipeline with Perl, R (R Core Team 2014), and BEDTools (Quinlan and Hall 2010). Perl was used to extract information from different datasets. R was used to build graphs to illustrate the repeat distribution in different genic regions, the identification of repetitive elements with respect to functional elements, GO term over-representation analysis and expression analysis of TEs. BED format file intersection was used to extract the overlapping regions between different datasets, with a lower limit of 25-bps. The UCSC Genome Browser (Dreszer, et al. 2012; Kent, et al. 2002) was used to download genome sequence data and genome annotations including RefSeq genes. RSEM (Trapnell, et al. 2012) was adopted to assemble RNA-Seq reads into transcripts and estimate their abundance (measured as transcripts per million (TPM)) Plots were generated using ggplot2 in R (Wickham 2009).

### Datasets

#### Genome And Annotations

NCBI’s Human genome and its annotation datasets (RefSeq hg19) (Pruitt, et al. 2007) were downloaded from the UCSC Genome Browser (Dreszer, et al. 2012; Ernst, et al. 2011). A total of 37,697 human RefSeq transcripts were merged into 18,777 genes by taking the longest transcript(s) that represented each distinct gene locus. Repetitive elements were downloaded from the RepeatMasker (http://www.repeatmasker.org) track of the UCSC Genome Browser. All repetitive sequence intervals were also de-duplicated to deal with potential overlapping repeat annotations. Overall, there were 5,298,130 human repetitive elements, which represented approximately 1.467Gb in the human genome.

#### Regulatory Element Datasets From Six Human Cell Lines

The regulatory element datasets from six human cell lines were downloaded from the UCSC Genome Browser. Each cell line dataset contained the annotation of six regulatory elements: 1) Active Promoters, 2) Weak Promoters, 3) Strong Enhancers, 4) Weak Enhancers, 5) Insulators and 6) Polycomb Repressed Regions. These regulatory element annotations were derived from different chromatin states that have been marked by histone methylation, acetylation and histone variants H2AZ, PolIII, and CTCF (Ernst, et al. 2011).

#### Gene Expression Datasets From Six Human Tissues

Human RNA-seq data from the Illumina bodyMap2 transcriptome (See Table S4 for detail) (http://www.ebi.ac.uk/ena/data/view/ERP000546) dataset was used to measure the association between TEs and the expression levels of genes containing TEs in six tissues.

#### The Distribution Of Repetitive Elements In The Human Genome

To assess how human TEs were distributed in genes, we compared different genic regions containing TEs. Based on Repbase (Jurka, et al. 2005; Kohany, et al. 2006) annotations identified by RepeatMasker (http://www.repeatmasker.org), repeat elements in human were divided into two categories: human-specific repeats, and repeats shared with different species. Human-specific repeats were those annotated with “Homo sapiens” or “primates” as their origin (See Table S8 for the list of human-specific repeat classes), whilst those remaining were categorized as shared repeats. Intergenic regions as well as the exons and introns within 5’UTR, CDS and 3’UTR regions from RefSeq genes (Meyer, et al. 2013) were then compared with these different categories of repeats. Next, we generated the summarised distributions of repetitive elements overlapping these regions by calculating the proportions of bases belonging to repetitive elements within each of the combined sets of regions, i.e.:

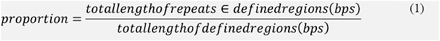

The code repository for the above can be found at https://github.com/UofABioinformaticsHub/RepeatElements.

#### The Occurrence Of Transposable Elements Within Regulatory Regions

We further explored the association between any TE and the regulatory elements defined above, by calculating the proportion of nucleotides within each of the five sets of genic regions (5’UTR, CDS-exon, CDS-intron, 3’UTR and Intergenic) that were part of a regulatory element for each of the six human cell lines. The proportion of nucleotides that were TEs within each regulatory element were also calculated for each genic region. All proportions were subsequently transformed using the logit function for model fitting across tissues (Table S1) using the model

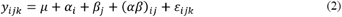

where *y_ijk_* is the logit transformed proportions representing **p(TE|RE)** across each genic region *i*, each regulatory element *j* and tissue *k,* such that *μ* is the overall mean, *α_i_* is the effect due to each genic region, *β_j_* is the effect due to each regulatory element with *(αβ)_ij_* representing any changes not accounted for in the first two terms. Tests for normality and homoscedasticity were performed using the Shapiro-Wilk test and Levene’s test respectively. Where violations of homoscedasticity were found robust standard errors were obtained using the sandwich estimator (Zeileis 2004). Confidence Intervals for pairwise comparisons were obtained as implemented in the R package *multcomp (Westfall 2008)* in order to control the Type I error at α=0.05 across the entire set of comparisons.

A specific TE analysis was then performed using six of the major human TE classes based on the Repbase classification system: *Alu*, L1, L2, MIR, LTR and DNA transposons. L1 elements were further resolved into either ancestral (L1M, L1PB and L1PA subfamilies) or recent/clade-specific (L1HS subfamily), based on their Repbase annotations. The proportions of nucleotides within each TE type that were also regulatory elements were calculated giving tissue-specific estimates of **p(TE|RE)**. Proportions were again transformed using the logit function, and the same analysis as above was performed.

#### Quality Control And Preprocessing Of The Gene Expression Data In Different Human Tissues

RNA-seq reads of six human tissues were first assessed using FastQC software (www.bioinformatics.babraham.ac.uk), to provide an overview of whether the raw RNA-Seq data contained any problems or biases before further analysis. Reads with poor-quality bases were trimmed (based on the results of FastQC with MINLEN set to 26) for subsequent data analysis. Table 4 showed the numbers of reads in raw RNA-Seq datasets and the statistics after the QC process by using Trimmomatic-0.32 (Bolger, et al. 2014). Then, we built transcript reference sequences using rsem-prepare-reference (Li and Dewey 2011) from Hg19 human RefSeq genes. The references were then input to rsem-calculate-expression (Li and Dewey 2011) using default parameters for all 6 tissues to obtain TPM based expression values.

Proximal promoter regions were defined as 1,000bp upstream of the gene transcription start sites based on the longest transcripts for each gene. *Alu*, MIR, L1, L2 and LTR repeat regions were then identified within the proximal promoters, 5’UTR, CDS and 3’UTR regions.

#### The Weighted Bootstrap Procedure For Assessing The Effects Of A TE In Each Genic Region

Many genes contain multiple transposable elements, with only a minority of genes containing a single TE (Figure S2). In order to assess any effects on transcription due to the presence of a single TE, a weighted bootstrap approach was devised. For a given TE within each genic region within each individual tissue, the frequencies of co-occurring TEs and combinations of TEs were noted. Uniform sampling probabilities were then used for the set of genes containing a specific TE in a specific region, whilst sampling weights were assigned to genes lacking the specific TE based on TE composition, such that the TE content of the sampled set of reference genes matched that of the test set of genes, based on the defined categories. Gene length was divided into 10 bins and these were included as an additional category when defining sampling weights. This ensured that two gene sets were obtained for each bootstrap iteration, which were matched in length and TE composition with the sole difference being the presence of the specific TE within each specific genic region (Figure S5). The mean difference in expression level, as measured by log(TPM), and the difference in the proportions of genes detected as expressed were then used as the variables of interest in the bootstrap procedure. The bootstrap was performed on sets of 1000 genes for 10,000 iterations using the proximal promoter as defined above, along with 5’UTR and 3’UTRs. When comparing expression levels, genes with zero read counts were omitted prior to bootstrapping. In order to compensate for multiple testing considerations, confidence intervals were obtained across the *m =* 90 tests at the level 1 – *α/m*, which is equivalent to the Bonferroni correction, giving confidence intervals, which controlled the FWER at the level *α =* 0.05. Approximate two-sided p-values were also calculated by finding the point at which each confidence interval crossed zero, and additional significance was determined by estimating the FDR on these sets of p-values using the Benjamini-Hochberg method.

#### Long Intergenic Non-Coding Rnas And Tes

Annotations for 8,196 previously described putative human lincRNAs were downloaded (Cabili, et al. 2011) and the distribution of TEs within regulatory elements in lincRNA exons and introns was obtained using the same methods as above. The previously described regression models were then used to analyse this dataset.

#### Association Of Functional Elements With Human Repetitive Elements

To demonstrate the potential functional significance of repetitive elements, the Database for Annotation, Visualization and Integrated Discovery (DAVID) (Dennis, et al. 2003) was used to perform the GO classification. We first extracted Gene-IDs from overlapping regions between different gene categories (1000bp proximal promoter, 5’UTR, 3’UTR, and the combination of these 3 regions) and TEs. These gene-lists were then submitted to the DAVID Functional Classification Tool. We chose the third level of GO terms to describe the over-represented functional terms for the three datasets and visualized the functional over-representation of overlapped genes using the R package *heatmap.2*. The p-value was applied in the GO analysis as the standard index to determine the degree of enrichment. The threshold for over-represented GO terms was set to an FDR (Benjamini-Hochberg method) less than 0.05. Protein-coding genes with Alus were also visualised with the UCSC genome browser (http://genome.ucsc.edu/) to compare their mRNA with various gene datasets and annotations.

#### Association Of Alternative Splicing And Protein Coding Regions Containing TEs

In order to assess the relationship between transposable elements and exonization, an alternative splicing annotation dataset (SIB Alt-splicing) was downloaded from the UCSC Genome Browser (http://genome.ucsc.edu/). These data were generated from RefSeq genes, Genbank RNAs and ESTs that aligned to the human genome. A total of 46,973 alternatively spliced transcripts were intersected with gene models containing transposable elements.

## ACKNOWLEDGEMENT

The authors wish to thank Dan Kortschak, Atma Ivancevic, Joy Raison, Reuben Buckley and Sim Lin Lim for valuable discussions and critical reading of drafts. This work was supported by grants from the National Natural Science Foundation of China (61272250 and 61472246) and the National Basic Research Program of China (2013CB956103). The funders had no role in study design, data collection and analysis, decision to publish, or preparation of the manuscript.

## AUTHOR CONTRIBUTION

DLA and CCW conceived, designed and managed the study. LZ collected the datasets, implemented the analysis pipeline, and analyzed the data. SMP analyzed the data. ZPQ, DFC, ZQH prepared datasets. LZ, DLA and CCW wrote and revised the manuscript. All authors reviewed and approved the final manuscript.

